# Driver-independent *lexAop-tdTomato.nls* reporter signal in the adult *Drosophila* proventriculus

**DOI:** 10.64898/2026.07.07.737111

**Authors:** Xinyue Zhou, Tianmu Zhang, Woo Jae Kim

## Abstract

Reporters are widely used in *Drosophila* genetics to visualize gene expression and cell lineages. However, uncharacterized limitations in specific reporter lines can lead to data misinterpretation. Here, we identify a consistent, driver-independent tdTomato signal in the adult proventriculus from the widely used *lexAop-tdTomato.nls* reporter line. This signal was observed across multiple *lexA* driver combinations and was directly detectable in *lexAop-tdTomato.nls* responder-alone adult proventriculi lacking any *lexA* driver and without antibody staining. In contrast, no comparable native red fluorescence was detected in larval proventriculi under the same no-antibody imaging condition. Mouse and rabbit anti-RFP immunostaining further supported the presence of proventriculus-associated tdTomato/RFP antigen in adult responder-alone animals. In larval responder-alone proventriculi, antibody-amplified staining was antibody-source-dependent: a detectable signal was observed only with rabbit anti-RFP, whereas mouse and rat anti-RFP produced no reliable detectable signal under the same staining condition. A driver-matched comparison using *lexAop-RFP.nls* did not reproduce the proventricular signal, arguing against detectable ectopic activity of the tested *lexA* driver in this tissue. However, because *lexAop-tdTomato.nls* and *lexAop-RFP.nls* differ in reporter/transgene architecture and possibly genomic insertion context, the underlying cause cannot be assigned specifically to the lexAop sequence. Our findings highlight the necessity of including driver-negative and no-antibody controls when using this reporter line in adult *Drosophila* proventriculus and gut studies.

## INTRODUCTION

The binary expression system, particularly the *GAL4/UAS* system, has been a cornerstone of genetic research in *Drosophila melanogaster*, enabling precise spatiotemporal control of gene expression(Fischer et al. 1988; Brand and Perrimon 1993). The complementary *lexA/lexAop* system offers an orthogonal tool for dual-labeling and intersectional studies(Lai and Lee 2006). Reporter lines such as *lexAop-tdTomato.nls* are widely employed for visualizing neuronal circuits and cell lineages due to the bright, stable fluorescence of tdTomato(Pfeiffer et al. 2008). However, the fidelity of transgenic reporters can be compromised by “leakage” or background expression, often resulting from enhancer activities at the genomic insertion site or basal promoter activity (Wong et al. 2002; Jory et al. 2012; Li et al. 2013). Such artifacts can lead to erroneous interpretation of expression patterns, especially in tissues with complex gene regulation like the gut. Here, we document a consistent driver-independent signal associated with the *lexAop-tdTomato.nls* reporter line in the adult proventriculus (PV). Because the *Drosophila* gut, including the foregut/proventriculus region, can present tissue-specific challenges for reporter interpretation, stringent driver-negative and reporter-matched controls are essential when evaluating apparent expression patterns in this tissue. By employing genetic crosses, responder-alone controls, native fluorescence imaging, antibody validation, and transcriptomic analyses, we characterize this phenomenon and propose critical control strategies.

## MATERIALS AND METHODS

### Fly Stocks and Husbandry

*Drosophila melanogaster* was raised on cornmeal-yeast medium at similar densities to yield adults with similar body sizes. *Drosophila* were kept in 12 h light: 12 h dark cycles (LD) at 25□ (ZT 0 is the beginning of the light phase, ZT12 beginning of the dark phase). The following stocks were used: *w1118; lexAop-tdTomato.nls* (Bloomington Stock Center, 66680), *w1118; FMRFa-lexA* (Bloomington Stock Center, 52824), *w1118; Lk*^*2A*^*-lexA* (Qidong Fungene, FBF00197), *w1118; Proc*^*2A*^*-lexA* (Bloomington Stock Center, 84432), *w; Tk*^*2A*^*-lexA* (Bloomington Stock Center, 84438), *w1118;; SIFaR*^*2A*^*-GAL4* (Bloomington Stock Center, 84689), *w1118; UAS-Stinger* (Bloomington Stock Center, 28864), *w1118; SIFaR*^*2A*^*-lexA* (Bloomington Stock Center, 84435), *w1118; lexAop-RFP.nls* (Bloomington Stock Center, 29956).

### Animal Ethics Statement

All experiments involving *Drosophila melanogaster* were conducted in accordance with the ARRIVE guidelines and complied with the U.K. Animals (Scientific Procedures) Act, 1986, EU Directive 2010/63/EU, and the National Research Council’s Guide for the Care and Use of Laboratory Animals.

### Tissue Dissection and Immunostaining

Adult guts were dissected from 5-day-old adult *Drosophila*, and larval proventriculi were dissected from third-instar larvae where indicated. Tissues were fixed in 4% formaldehyde at room temperature for 30 minutes. The sample was then washed three times (5 minutes each) in 1% PBT and then blocked in 5% normal goat serum for 30 minutes. Subsequently, the sample was incubated overnight at 4□ with primary antibodies in 1% PBT, followed by the addition of fluorophore-conjugated secondary antibodies for one hour at room temperature. Antibodies were used at the following dilutions: rabbit anti-RFP (1:500, 1:1000, Rockland, 600-401-379); mouse anti-RFP (1:500, 1:1000, Thermo, MA5-15257); rat anti-RFP (1:500, 1:1000, ChromoTek, 5F8); chicken anti-GFP (1:500, Invitrogen, A10262), Alexa-488 donkey anti-chicken (1:200, Jackson ImmunoResearch), Alexa-555 goat anti-rabbit (1:200, Invitrogen), Alexa-555 donkey anti-rat (1:200, Invitrogen, A48270), Alexa-555 goat anti-mouse (1:200, Invitrogen, A32727).

### Microscopy imaging and image processing

Confocal images were acquired using a Zeiss LSM 880 microscope with consistent laser power and detector gain settings within each comparative experiment. Image processing (maximum intensity projection, brightness/contrast adjustment applied equally across compared images) was performed using Fiji/ImageJ software (Schindelin et al. 2012).

### Single-nucleus RNA-sequencing analyses

The single-nucleus RNA-seq dataset analyzed in this study was obtained from the Fly Cell Atlas (Li et al. 2022) and visualized using the Fly SCope website (https://scope.aertslab.org/#/FlyCellAtlas//welcome). Raw and processed datasets are available through the Fly Cell Atlas resources, including SCope (https://flycellatlas.org/scope) and ASAP (https://asap.epfl.ch/fca). For t-SNE visualization, we selected the relevant tissue sessions from the “10X Cross-tissue” dataset. The following display settings were used: “Log transform,” “CPM normalize,” “Expression-based plotting,” “Show labels,” and “Dissociate viewers” were enabled, and both “Point size” and “Point alpha level” were set to maximum. Gene coexpression was visualized using the color scheme provided by Fly SCope, with each color corresponding to the genes selected on the left, right, and bottom panels of the viewer. Yellow, cyan, purple, and white signals indicate overlap between red/green, green/blue, red/blue, and all three selected markers, respectively. The same visualization settings were applied consistently across all t-SNE plots.

## RESULTS

### Consistent, driver-independent tdTomato signal in the adult proventriculus

To test whether *lexAop-tdTomato.nls* faithfully reports driver activity, we crossed it with five independent *lexA* drivers, including both traditional enhancer-trap and T2A knock-in lines (*FMRFa-lexA, Lk*^*2A*^*-lexA, Proc*^*2A*^*-lexA, Tk*^*2A*^*-lexA*, and *SIFaR*^*2A*^*-lexA*). These drivers are known to be expressed exclusively in specific CNS neuronal populations, with no documented activity in the adult PV or other gut regions (Nässel and Winther 2010; Leader et al. 2017). Consistently, FlyAtlas2 RNA-seq data show negligible transcript levels of these genes in the PV/cardia, and a previous review reported that *tachykinin* (*Tk*) is not expressed in the *Drosophila* foregut(Nässel 2018) .

To validate this at single-cell resolution, we performed co-localization analysis using the PV marker genes *CG43673* and *Mur29B*(Zhu et al. 2024) . As shown in Fig. S1, no significant overlap was observed between these PV markers and *Proc, Lk, Tk*, or *FMRFa*, supporting the conclusion that these neuropeptide genes are not detectably expressed in PV epithelial cells under this dataset. These results further support that the tdTomato signal observed in the PV is unlikely to originate from endogenous driver activity.

Nevertheless, all five crosses produced an anatomically consistent tdTomato signal in the adult PV (Fig. 1A–E). This cross-driver consistency supports the interpretation that the PV signal is driver-independent and associated with the *lexAop-tdTomato.nls* reporter line rather than genuine activity of any single *lexA* driver. To exclude the possibility that the observed signal reflects endogenous *SIFaR* expression, we performed a parallel control using *SIFaR*^*2A*^*-GAL4* — a knock-in driver generated *via* the same T2A-fusion strategy (Diao and White 2012) — to drive *UAS-Stinger* (nuclear GFP). No GFP signal was detected in the PV (Fig. 1F), supporting the conclusion that the PV signal observed with *SIFaR*^*2A*^*-lexA; lexAop-tdTomato.nls* is not explained by endogenous SIFaR expression or by detectable ectopic activity of the *SIFaR*^*2A*^ driver in this tissue.

**Figure 1.**
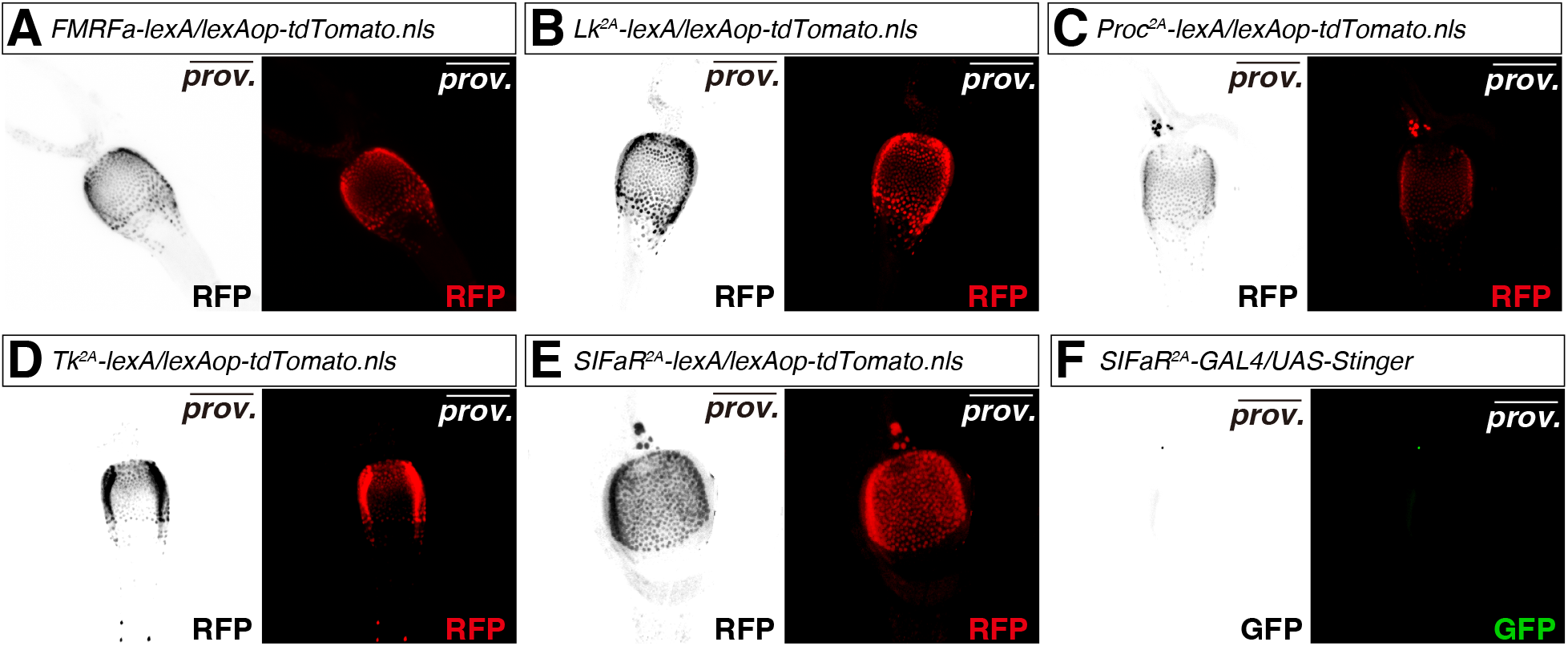
The *lexAop-tdTomato.nls* reporter exhibits a driver-independent signal in the adult proventriculus. (A–E) Confocal images of adult *Drosophila* proventriculus showing tdTomato fluorescence after crossing *lexAop-tdTomato.nls* with five different *lexA* drivers: (A) *FMRFa-lexA*, (B) *Lk*^*2A*^*-lexA*, (C) *Proc*^*2A*^*-lexA*, (D) *Tk*^*2A*^*-lexA*, (E) *SIFaR*^*2A*^*-lexA*. All crosses consistently showed tdTomato signal in the proventriculus region. (F) Negative control using *SIFaR*^*2A*^*-GAL4* driving *UAS-Stinger* (GFP). No specific GFP signal was detected in the proventriculus, supporting the conclusion that the PV signal is not explained by endogenous SIFaR expression or detectable ectopic activity of the *SIFaR*^*2A*^ driver . Scale bar: 50 µm.

### Native fluorescence and anti-RFP controls distinguish adult and larval proventriculus-associated reporter signals

To directly determine whether the reporter-associated signal is detectable without antibody amplification, we imaged *lexAop-tdTomato.nls* responder-alone proventriculi lacking any *lexA* driver under no-antibody conditions. Native red fluorescence was detected in the adult proventriculus (Fig. S2D). In contrast, no comparable red fluorescence was observed in larval proventriculi processed and imaged under the same no-antibody condition (Fig. S2E). These results indicate that the native reporter-associated red fluorescence is detectable in the adult proventriculus but not in the larval proventriculus under the no-antibody imaging condition tested. Anti-RFP immunostaining further supported the presence of tdTomato/RFP antigen in the adult proventriculus. Mouse and rabbit anti-RFP antibodies produced clear proventriculus-associated staining in *lexAop-tdTomato.nls* responder-alone adults lacking any *lexA* driver, whereas the rat anti-RFP antibody produced weak or undetectable staining under our conditions (Fig. 2A–F). We therefore base the adult antibody-validation conclusion primarily on the reproducible native tdTomato fluorescence together with mouse and rabbit anti-RFP staining.

**Figure 2.**
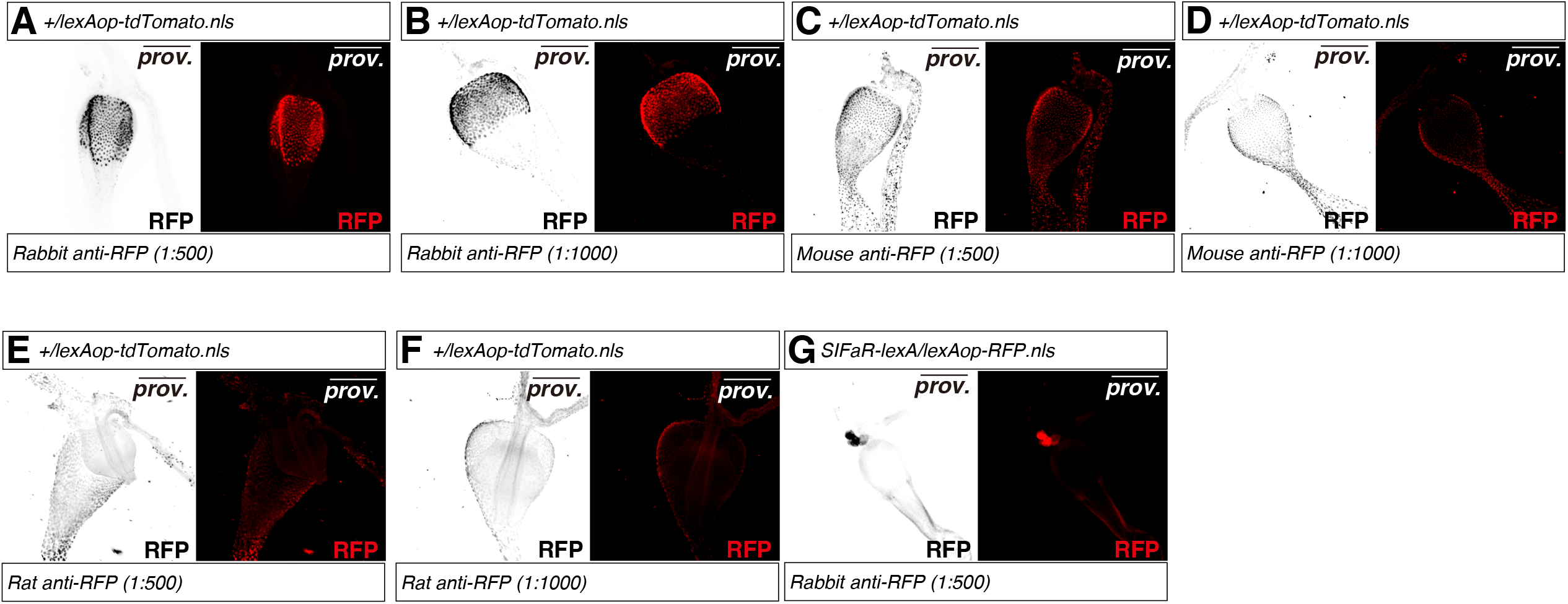
Anti-RFP validation of the adult proventriculus-associated signal in *lexAop-tdTomato.nls* responder-alone animals. Immunostaining was performed on adult proventriculus tissue from *lexAop-tdTomato.nls* homozygotes without any *lexA* driver. (A, B) Staining results using rabbit anti-RFP antibody at (A) 1:500 and (B) 1:1000 dilutions. (C, D) Staining results using mouse anti-RFP antibody at (C) 1:500 and (D) 1:1000 dilutions. (E, F) Staining results using rat anti-RFP antibody at (E) 1:500 and (F) 1:1000 dilutions. (G) Driver-matched control using *SIFaR*^*2A*^*-lexA* driving *lexAop-RFP.nls*. No comparable RFP signal was detected in the adult proventriculus under the same driver condition, arguing against detectable ectopic activity of *SIFaR*^*2A*^*-lexA* in this tissue. Mouse and rabbit anti-RFP antibodies produced clear proventriculus-associated staining, whereas rat anti-RFP staining was weak or less reliable under our conditions. Scale bar: 50 µm.

We also examined larval *lexAop-tdTomato.nls* responder-alone proventriculi after anti-RFP immunostaining at 1:500 dilution with DAPI nuclear counterstaining. In contrast to the no-antibody condition, antibody-amplified staining showed antibody-source-dependent differences. Rabbit anti-RFP produced a detectable proventriculus-associated signal, whereas mouse and rat anti-RFP produced no reliable detectable signal under the same staining condition (Fig. S2A–C). These results indicate that the absence of native red fluorescence in larvae should be distinguished from antibody-amplified staining outcomes, and that the larval anti-RFP signal is detectable only with the rabbit anti-RFP antibody under the conditions tested.

Because the reporter contains a nuclear localization signal, we performed DAPI nuclear counterstaining in the indicated adult and larval proventriculus controls. DAPI staining provided a nuclear reference for the proventricular tissue and helped distinguish the red-channel signal from tissue morphology. However, we do not use DAPI staining as quantitative evidence for subcellular colocalization. We therefore conservatively describe the phenotype as a proventriculus-associated tdTomato/RFP signal.

To further distinguish driver-dependent expression from reporter-line-associated signal, we performed a driver-matched comparison using an independent *lexA*-responsive reporter, *lexAop-RFP.nls. SIFaR*^*2A*^*-lexA* was crossed in parallel to *lexAop-tdTomato.nls* and *lexAop-RFP.nls*. While *SIFaR*^*2A*^*-lexA; lexAop-tdTomato.nls* showed a strong proventricular signal (Fig. 1E), no comparable signal was detected with *SIFaR2A-lexA; lexAop-RFP.nls* under the same driver condition (Fig. 2G). This result argues against detectable ectopic *SIFaR*^*2A*^*-lexA* activity in the adult proventriculus.

However, because *lexAop-tdTomato.nls* and *lexAop-RFP.nls* are not identical reporter constructs and differ in reporter sequence, regulatory architecture, and/or genomic insertion context, this comparison does not identify the precise causal element responsible for the *lexAop-tdTomato.nls*-associated signal.

This differential outcome indicates that the PV signal is associated with the *lexAop-tdTomato.nls* reporter line under the conditions tested, but does not distinguish between effects of the *13XLexAop2-IVS*-containing transgene architecture, the tdTomato reporter cassette, and the genomic insertion site.

## DISCUSSION

This study identifies a consistent, driver-independent signal associated with the *lexAop-tdTomato.nls* reporter line in the adult *Drosophila* proventriculus. The key evidence is the detection of native red fluorescence in adult *lexAop-tdTomato.nls* responder-alone proventriculi lacking any *lexA* driver and without antibody staining (Fig. S2D). In contrast, no comparable native red fluorescence was detected in larval proventriculi under the same no-antibody condition (Fig. S2E), suggesting that the native reporter-associated red fluorescence is enriched in the adult proventriculus under the conditions tested. The same anatomical signal was also observed across multiple independent *lexA* driver combinations (Fig. 1A–E), supporting the interpretation that the signal is not caused by genuine activity of any single *lexA* driver. Anti-RFP immunostaining with mouse and rabbit antibodies further supported the presence of tdTomato/RFP antigen in the adult proventriculus (Fig. 2A–D), whereas rat anti-RFP staining was weak or undetectable under our conditions (Fig. 2E, F). In larval proventriculi, anti-RFP staining at 1:500 showed antibody-source-dependent differences: rabbit anti-RFP produced a detectable proventriculus-associated signal, whereas mouse and rat anti-RFP produced no reliable detectable signal under the same condition (Fig. S2). These observations emphasize that native fluorescence and antibody-amplified staining should be interpreted separately. The experimental results are also supported by Fly Cell Atlas single-nucleus RNA-seq analyses, which show little or no overlap between proventriculus marker genes and the tested driver-associated endogenous neuropeptide genes (Fig. S1). Together, these results indicate that the observed adult proventricular signal should not be interpreted as bona fide *lexA* driver activity without appropriate driver-negative and no-antibody controls.

The mechanism underlying this reporter-line-associated signal remains unresolved. One possibility is a genomic position effect, in which the insertion site of the *lexAop-tdTomato.nls* transgene places the reporter in a chromatin environment or near regulatory elements active in the adult proventriculus. A second possibility is that sequence elements within the *lexAop-tdTomato.nls* transgene itself, including the *13XLexAop2-IVS* regulatory architecture, vector backbone, minimal promoter, or tdTomato reporter cassette, contribute to low-level proventriculus activity. The absence of a comparable signal in *SIFaR2A-lexA; lexAop-RFP.nls animals* (Fig. 2G) supports the conclusion that the tested driver does not show detectable proventricular activity, but it does not identify the causal element responsible for the *lexAop-tdTomato.nls*-associated signal because *lexAop-tdTomato.nls* and *lexAop-RFP.nls* are not identical reporter constructs. Because we were not able to test an additional matched *13XLexAop2-IVS* reporter inserted at an independent genomic site, our data cannot distinguish between a genomic position effect and transgene-intrinsic activity. Future comparisons using matched *13XLexAop2-IVS* reporter constructs at different genomic sites, or independent insertions at the same genomic landing site, will be required to determine the causal mechanism.

Although *lexAop-tdTomato.nls* contains a nuclear localization signal, our DAPI-stained controls are used here as qualitative nuclear references rather than as quantitative colocalization evidence. Therefore, we conservatively describe the phenotype as a proventriculus-associated tdTomato/RFP signal instead of making a strong claim about precise subcellular localization.

## ACKNOWLEDGEMENTS

We sincerely thank Professor Woo Jae Kim for his invaluable guidance, insightful discussions, and continuous support throughout this study. This research was supported by Startup funds from HIT Center for Life Science (HCLS) to WJK.

## DECLARATIONS

### CONSENT TO PARTICIPATE

The research described in this paper does not involve any human participants. Therefore, no consent to participate was obtained.

### CONSENT FOR PUBLICATION

All authors have given consent for the publication of this work.

### AVAILABILITY OF DATA AND MATERIALS

All data and reagents reported in this paper will be shared by the corresponding author upon request.

### COMPETING INTERESTS

The authors declare no competing interests

### AUTHORS’ CONTRIBUTIONS

**Conceptualization:** Woo Jae Kim.

**Data curation:** Xinyue Zhou, Tianmu Zhang, Woo Jae Kim.

**Formal analysis:** Xinyue Zhou, Tianmu Zhang, Woo Jae Kim.

**Funding acquisition:** Woo Jae Kim.

**Investigation:** Woo Jae Kim.

**Methodology:** Woo Jae Kim.

**Project administration:** Woo Jae Kim.

**Resources:** Woo Jae Kim.

**Supervision:** Woo Jae Kim.

**Validation:** Tianmu Zhang, Woo Jae Kim.

**Visualization:** Xinyue Zhou, Tianmu Zhang, Woo Jae Kim.

**Writing – original draft:** Xinyue Zhou, Tianmu Zhang.

**Writing – review & editing:** Xinyue Zhou, Tianmu Zhang.

## DECLARATION OF GENERATIVE AI AND AI-ASSISTED TECHNOLOGIES IN THE WRITING PROCESS

During the creation of this work, the author(s) utilized DeepSeek and Perplexity to rephrase English sentences, verify English grammar, and detect plagiarism, as none of the authors of this paper are native English speakers. After using this tool/service, the author(s) reviewed and edited the content as needed and take(s) full responsibility for the content of the publication.

## DATA AVAILABILITY STATEMENT

Strains are available upon request. The authors affirm that all data necessary for confirming the conclusions of the article are present within the article and figures.

## FIGURE LEGENDS

**Figure S1.**
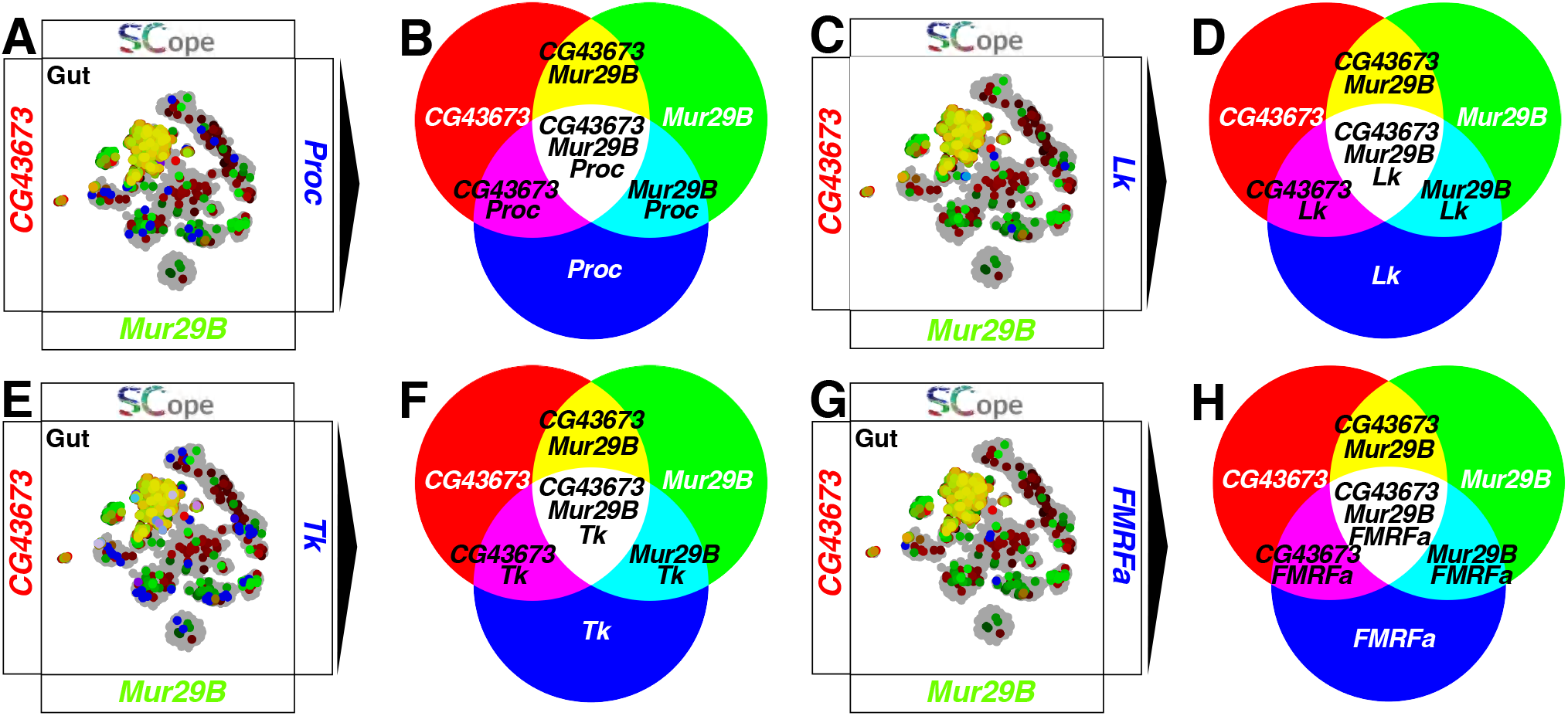
Single-cell RNA-seq analysis confirms absence of neuropeptide gene expression in proventricular epithelial cells. (A-H) SCOPE scRNA-seq datasets reveal tissues colored by expression of proventriculus marker genes *CG43673* (red) and *Mur29B* (green) with (blue) (A-B) *Proc*, (C-D) *Lk*, (E-F) *Tk*, (G-H) *FMRFa*.

**Figure S2.**
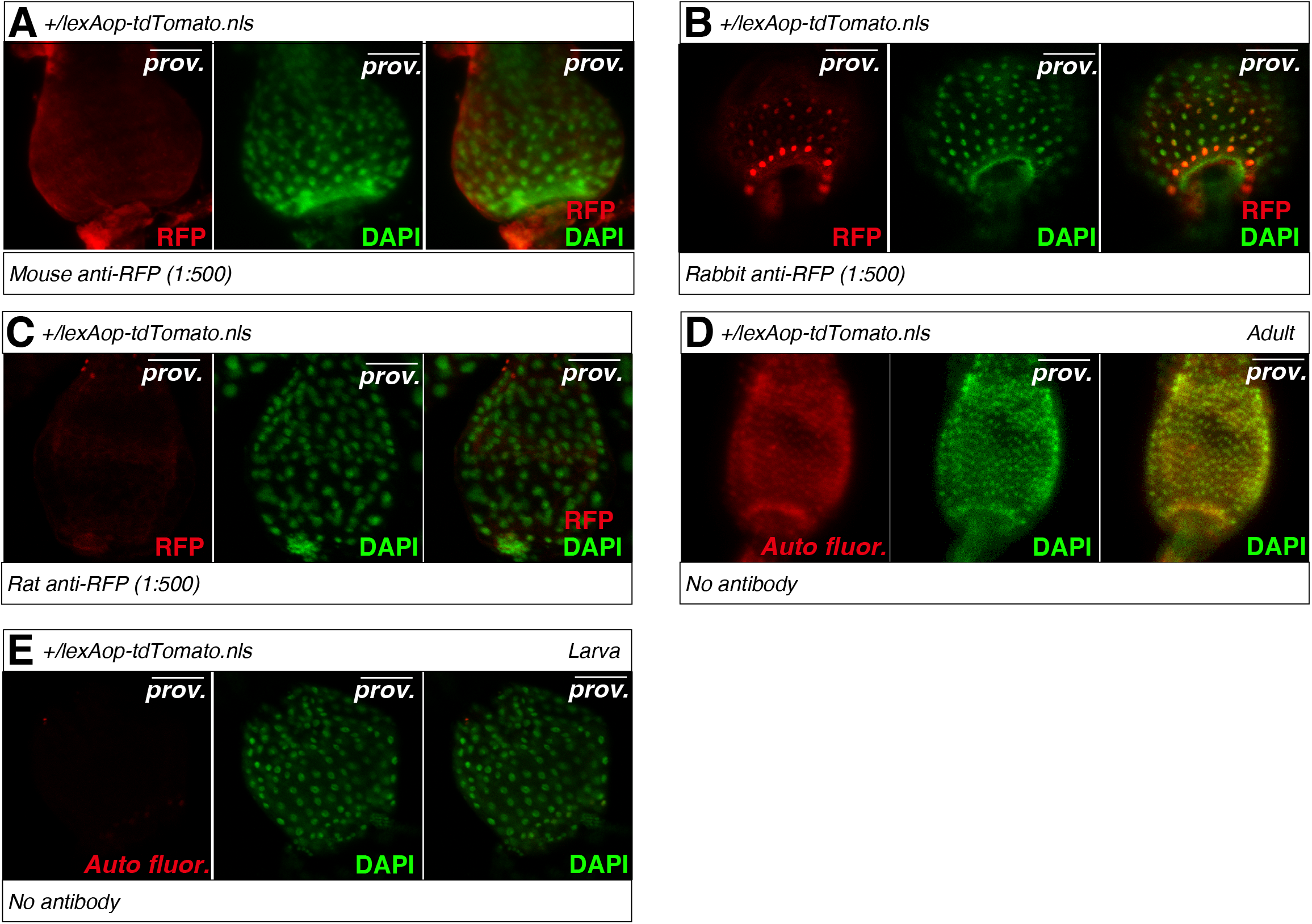
Larval anti-RFP staining and no-antibody proventriculus controls in *lexAop-tdTomato.nls* responder-alone animals. (A) Larval proventriculi from third-instar *lexAop-tdTomato.nls* responder-alone animals lacking any *lexA* driver were stained with anti-RFP antibodies at 1:500 dilution. DAPI was used as a nuclear counterstain and is shown in green. (A) Mouse anti-RFP staining produced no reliable detectable proventriculus-associated RFP signal under the condition tested. (B) Rabbit anti-RFP staining produced a detectable proventriculus-associated RFP signal. (C) Rat anti-RFP staining produced no detectable proventriculus-associated RFP signal under the same staining condition. (D) Adult proventriculus from *lexAop-tdTomato.nls* responder-alone animals imaged without antibody staining. A red-channel signal was detected in the adult proventriculus. (E) Third-instar larval proventriculus from *lexAop-tdTomato.nls* responder-alone animals imaged without antibody staining. No comparable red-channel signal was detected in the larval proventriculus. Scale bar: 50 µm.

## Notes

### Competing Interest Statement

The authors have declared no competing interest.

